# Deletion of OTUD7B in astrocytes protects against cerebral malaria by inhibiting microvesicle-induced TRAF3/TRAF6-mediated neuroinflammation

**DOI:** 10.64898/2026.04.16.717638

**Authors:** Kunjan Harit, Jakob Johann Schmidt, Ruth Beckervordersandforth, Dirk Schlüter, Gopala Nishanth

**Author notes:** Correspondence: Gopala Nishanth, Institute of Medical Microbiology and Hospital Epidemiology, Hannover Medical School, Carl-Neuberg-Straße 1, 30625 Hannover; Germany. Equal contribution (Dirk Schlüter, Gopala Nishanth).

## Abstract

Cerebral malaria is a severe neurological complication of *Plasmodium falciparum* infection. Damage of the blood-brain barrier (BBB) and endothelial dysfunction are established drivers of the disease pathology, however, whether astrocytes, a major constituent of the BBB, influence the disease outcome remains unclear. Using the murine model of experimental cerebral malaria (ECM), we show that astrocytes decisively regulate the outcome of ECM and the deubiquitinating enzyme OTUD7B in astrocytes fosters the disease. Mice lacking astrocytic OTUD7B showed reduced brain pathology and were protected from ECM compared with wildtype littermate controls. Transcriptomic profiling of ex vivo–isolated astrocytes revealed reduced proinflammatory chemokines and cytokines in the absence of OTUD7B. Plasmodium infection–associated microvesicles triggered a pro-inflammatory response in astrocytes, which was dependent on OTUD7B. Mechanistically, OTUD7B cleaved K48-linked ubiquitin chains from TRAF3 and TRAF6 upon stimulation with microvesicles or activation of TLR3/TLR9 by plasmodial nucleic acids. The OTUD7B-dependent TRAF3 and TRAF6 stabilization led to sustained NF-κB and p38 MAP kinase signaling and CXCL10 expression. Therapeutic silencing of CNS Otud7b or Cxcl10 expression after disease onset protected mice from ECM, identifying the cerebral OTUD7B-Cxcl10 axis as an attractive therapeutic target.

## Introduction

Cerebral malaria (CM) is a life-threatening neurological complication of *Plasmodium falciparum* infection and a major cause of mortality, particularly in children living in sub–Saharan Africa and adults in Asia (World Malaria Report 2025, ^1^). Post-mortem studies identified blood-brain barrier (BBB) dysfunction, brain endothelial activation, neurovascular leukocyte accumulation, cerebral edema, and brain hemorrhages, as the hallmarks of CM ^2–4^. Multiple therapies targeting downstream inflammatory mediators including corticosteroids, anticoagulants, and TNF neutralization have failed to improve the outcome of CM suggesting that the early upstream regulators of the neuroinflammation still remain to be identified ^5–7^.

Experimental cerebral malaria (ECM), induced by *Plasmodium berghei* ANKA (PbA) infection recapitulates many features of the human disease and is a widely used model to study human CM ^8–10^. The key factors associated with ECM pathogenesis include (i) sequestration of parasitized RBCs in the brain vasculature, (ii) intravascular coagulation, (iii) activation and dysfunction of endothelial cells, and (iv) a pronounced neuroinflammatory immune response. All of these processes take place within the neurovascular unit, where the endothelial cells and brain-resident parenchymal cells respond to the pathogenic cues, regulate local inflammation and thereby determine the vascular integrity ^11–13^.

Astrocytes are a key component of the neurovascular unit which provide physical support to the BBB via their glia limitans and maintain proper functioning of the brain through regulation of nutrients delivery and transport of ions and neurotransmitters ^14,15^. In addition to these homeostatic functions, astrocytes can also shape neuroinflammation via chemokine- and cytokine- mediated recruitment of leukocyte and, thereby, control the immune response in the CNS ^16^. Medana et al., 2001, observed altered astrocyte distribution and degeneration of astrocytes at the peak stage of ECM suggesting that astrocytes might play an important role in the pathogenesis of CM ^17^. In concurrence, we and others have shown that astrocytes are activated during ECM ^18–20^.

In addition to pro-inflammatory cytokines, circulating *Plasmodium*-infected red blood cells-and small extracellular vesicles (EVs)/microvesicles derived from parasitized red blood cells, endothelial cells, platelets and leukocytes contribute to BBB damage in ECM. Microvesicles are rich in phosphatidylserine, and their cargo comprises of immunomodulatory proteins, lipids, and nucleic acids ^21–23^. However, whether astrocytes can sense plasmodium infection-associated microvesicles and whether this impacts on the outcome of ECM is unresolved.

Physiological and pathological cellular reactions are critically determined by the reversible process of ubiquitination and deubiquitination, but their function in cerebral malaria is largely unknown. Ubiquitination is a process in which small regulatory proteins known as ubiquitin are covalently linked to the substrate protein through one of its seven lysine residues (K6, K11, K27, K29, K33, K48, K63) or the N-terminal methionine (M1). Depending upon the type of ubiquitin linkage the fate of the target substrate is determined ^24,25^. The best-characterized are K48 and K11-linked polyubiquitination, which lead to proteasomal degradation and K63-linked polyubiquitination, which determine nondegradative function such as protein-protein interaction and signal transduction ^24,25^. The process of ubiquitination can be reversed by deubiquitinating enzymes (DUBs) which cleave ubiquitin chains from the substrate. OTUD7B, also known as Cezanne, is a DUB with a preference for K11, K48 and K63-linked ubiquitin chains ^26–28^.

OTUD7B plays a pivotal role in the regulation of diverse inflammatory signaling pathways. Importantly, the effects of OTUD7B on immune signaling are highly cell-type specific. In endothelial cells, OTUD7B suppresses TNF-induced NF-κB signaling by removing K63-linked ubiquitin chains from RIPK1 ^29^, whereas in T cells OTUD7B impairs T cell receptor signaling by deubiquitinating ZAP70 ^30^. With respect to the CNS, we identified that in ECM, OTUD7B protects against TNF-driven apoptosis of dendritic cells by deubiquitinating the E3 ubiquitin ligase TRAF2 in ECM ^31^. In a model of CNS autoimmunity, we recently demonstrated that astrocyte-specific OTUD7B limits and attenuates neuroinflammation by inhibiting degradation of glial fibrillary acidic protein (GFAP) and preventing TNF-induced astrocyte hyperinflammation by sequential K63- and K48- deubiquitination of RIPK1^32^. These studies show that the function of OTUD7B in infection and inflammation is cell type-specific and may depend on the inflammatory milieu.

To address whether (i) astrocytes play an important role in ECM, (ii) OTUD7B regulates microvesicle-mediated astrocyte activation and (iii) astrocytic OTUD7B impacts on the pathogenesis of ECM, we studied ECM in mice lacking *Otud7b* selectively in astrocytes (GFAP-Cre OTUD7b^fl/fl^) ^32^. Here, we identified astrocytes as critical regulators of ECM and uncover an unexpected, pro-inflammatory role of OTUD7B in Plasmodium-driven CNS inflammation. OTUD7B fostered microvesicle-induced NF-κB and p38 MAP kinase signaling in astrocytes by K48-deubiquitination of TRAF3 and TRAF6. This augmented production of the chemokine CXCL10, a key factor recruiting pathogenetic CD8^+^ T cells to the brain. Importantly, therapeutic inhibition of Otud7b and Cxcl10, respectively, after disease onset preserved BBB integrity and rescued mice from severe ECM.

## Materials and Methods

### Mice

Otud7b^fl/fl^ mice were crossed with GFAP-Cre mice to generate astrocyte-specific OTUD7B-deficient mice GFAP-Cre Otud7b^fl/fl^ as previously described by us ^32^. Littermate Otud7b^fl/fl^ mice were used as controls. The mice were genotyped with primers specific for floxed *Otud7b* and GFAP-cre ^32^. The animals were kept under specific pathogen-free (SPF) conditions in the animal facilities of the Otto-von-Guericke University Magdeburg (Magdeburg, Germany) and Hannover Medical School (Hannover, Germany). Both male and female mice aged 8–10 weeks were used. The animal experiments were carried out in accordance with the European animal protection law and approved by local authorities (Landesverwaltungsamt Halle, file number 42502-2-1260).

### Experimental cerebral malaria model

*Plasmodium berghei* ANKA (PbA) was used for mouse infection experiments. Parasite stocks were generated by infecting C57BL/6 mice with infected red blood cells (iRBCs). On day 7, postinfection, blood was collected and mixed with Alsever’s solution supplemented with 10% glycerol. Mice were injected intraperitoneally with 1 × 10⁶ *Plasmodium berghei* ANKA–infected red blood cells (iRBCs).

### Assessment of parasitemia

Thin blood smears were prepared from tail blood, fixed in methanol, and stained with 10% Giemsa solution diluted in PBS. The parasite load was determined by calculating the percentage of infected red blood cells across 10 randomized light-microscopy fields.

### Evans Blue staining

On day 7 p.i., mice were intravenously (i.v.) injected with 100 μl of 1% Evans Blue dye diluted in PBS. After 1h, mice were anesthetized with ketamine and xylazine and perfused intracardially with 0.9% NaCl. Perfused brains were collected and photographed.

### Histology and immunostaining

Mice anesthetized with isoflurane were transcardially perfused with 0.1 M phosphate-buffered saline (PBS), followed by 4% paraformaldehyde (PFA) in PBS. After paraffin embedding the brain tissue was cut into thin sections of 5 µm and used for hematoxylin and eosin staining and immunostaining with antibodies against: rabbit anti-GFAP (1:1000), rabbit anti CD31 (1:1000), and rabbit anti CD8 (1:1000), The tissue sections were subsequently blocked with appropriate sera (1:10 in PBS) according to the source of the primary antibody and incubated with the respective primary antibodies at room temperature. After washing, the secondary antibody was added for 1 h and was detected using an avidin–biotin complex (ABC) method with 3,3′-diaminobenzidine (DAB) and H₂O₂ as the chromogenic substrate.

Immunofluorescence staining was performed as described earlier ^32^, paraffin tissue sections were deparaffinized by incubation at 60 °C for 1 h, followed by two 15-min washes in xylene to remove residual paraffin. The tissue was subsequently rehydrated and incubated in 0.5% sodium borohydride (NaBH₄) for 30 min at room temperature to reduce autofluorescence. Antigen retrieval was performed by incubating sections in 10 mM citric acid buffer containing 2 mM EDTA and 0.05% Tween-20 for 15 min at 95 °C, followed by cooling for 20 min at room temperature.

After washing in PBS containing 1% Triton X-100 (PBS-T) for 30 min and PBS for 10 min, sections were incubated at 4 °C with primary antibodies diluted in blocking solution (3% normal donkey serum, 0.5% Triton X-100). The following primary antibodies were used: rabbit anti-GFAP (1:1000), rabbit anti-OTUD7B (1:1000), mouse anti-Sox2 (1:500), and goat anti-SOX9 (1:500). Following two 15-min washes in PBS, sections were incubated overnight at 4 °C with secondary antibodies diluted in blocking solution. Secondary antibodies included Alexa Fluor 488–conjugated donkey anti-goat (1:400), Alexa Fluor 488–conjugated donkey anti-mouse (1:400), Cy3-conjugated donkey anti-rat (1:400), and Cy5-conjugated donkey anti-rabbit (1:400).

After removal of secondary antibodies, nuclei were counterstained with DAPI (1:10,000). Sections were washed three times for 10 min in PBS, mounted with Aquapolymount, and stored at 4 °C. Images were acquired using a Zeiss Axio Observer 7 inverted microscope equipped with an ApoTome.2, an Axiocam 503 camera, and a Colibri 7 LED light source, as well as a Zeiss LSM 780 confocal microscope with 405, 488, 559, and 633 nm lasers and ×20, ×40, and ×63 objective lenses. ApoTome image acquisition was performed using ZEN 2.6 Pro software.

### Primary astrocyte cultures and treatment

Primary astrocytes were isolated from postnatal day 0–1 (P0–P1) mice and cultured in DMEM containing 1% glutamine, 10% FCS, and 1% penicillin/streptomycin. The purity of astrocyte cultures was more than 95%, as assessed by flow cytometry with antibodies against CD11b and ACSA-2. Serum-derived microvesicles were isolated by differential centrifugation as described below (or move it above?). For stimulation experiments, astrocytes were either treated with purified serum microvesicles at a concentration corresponding to 1× 10⁸ vesicles/ ml or were stimulated with TLR3 agonist Polyinosinic–polycytidylic acid (Poly I:C; 10 µg/ml) or the TLR9 agonist ODN1585 (10 µg/ml). Untreated cells or vehicle-treated cells served as controls.

### Isolation of microvesicles

Whole blood was collected by cardiac puncture of PbA infected C57BL/6 mice at day 7 p.i. Serum was obtained by centrifugation at 2,000 × g for 10 min at 4 °C, followed by a second centrifugation at 2,000 × g for 10 min to remove residual cells. Platelets and cellular debris were depleted by centrifugation at 10,000 × g for 30 min at 4 °C. The supernatant was then centrifuged at 20,000 × g for 60 min at 4 °C to pellet microvesicles, which were washed once in sterile, 0.22-µm–filtered PBS and centrifuged again at 20,000 × g for 60 min. Purified microvesicles were resuspended in PBS.

Microvesicles were identified by Annexin V staining and analyzed by flow cytometry. Size distribution was assessed by co-analysis with sterile polystyrene size-calibration beads, and vesicles were gated within the expected microvesicle size range of approximately 100–1,000 nm.

### In vitro siRNA treatment

For siRNA-mediated knockdown of TRAF3 and TRAF6 in vitro, primary astrocytes were transfected with 5 µM of TRAF3 and TRAF6-specific siRNA for 48h according to the manufacturer’s instructions. Thereafter, the cells were stimulated with serum derived microvesicles.

### In vivo siRNA knockdown

For in vivo gene silencing, siRNA targeting *Otud7b* or *Cxcl10*, as well as a scrambled control siRNA, was administered intranasally on day 5 p.i. at a concentration of 5µg per mouse. siRNAs were prepared in sterile, Nuclease-free water and delivered dropwise into the nostrils of mice to facilitate nasal uptake. The mice were maintained in a supine position for several minutes following administration to ensure efficient delivery. Knockdown efficiency was assessed at the indicated time points by quantitative PCR or by WB from the brain tissue.

### Western blot

Samples from primary astrocytes and mouse organs were lysed on ice in RIPA lysis buffer supplemented with PhosSTOP, phenylmethylsulfonyl fluoride (PSMF) and protease inhibitor cocktail. Cell lysates were pre-cleared by centrifugation at 14,000 rpm for 15mins at 4°C. Supernatant was collected and quantified by BCA assay according to the manufacturer’s protocol. Protein samples were heated in lane marker reducing sample buffer at 99°C for 5 min. Equal amounts of samples were separated by SDS-PAGE and subsequently transferred to polyvinylidene difluoride (PVDF) membranes, which were blocked with 5% BSA at room temperature for 1h, followed by incubation with the mentioned primary antibodies at 4°C overnight.

Blots were developed using the ECL Plus Kit and images were captured on an Intas Chemo Cam Luminescent Image Analysis system (INTAS). Quantification and analysis of WB images was performed with the LabImage 1D software.

### Co-immunoprecipitation (Co-IP)

Whole cell lysates from astrocytes were precleared by incubation with GammaBind G Sepharose beads with gentle shaking at 4°C for 2 h. After removal of beads by centrifugation, samples were incubated with specific antibodies under continuous shaking at 4°C overnight. The following day, antibody-protein complexes were captured by incubating samples with GammaBind G Sepharose beads at 4°C for 2 hours. Thereafter, the beads were washed with ice cold PBS thrice, resuspended in 2x lane marker reducing sample buffer and boiled at 99°C for 5 min. Samples were centrifuged and supernatant was collected for WB analysis.

### Quantitative RT-PCR

Total mRNA was isolated from brain tissue or astrocytes in buffer RLT using the RNeasy Mini Kit according to the manufacturer’s protocol. mRNA was reverse transcribed into cDNA with the SuperScript Reverse Transcriptase Kit. Quantitative RT-PCR was performed with a LightCycler 480 system using TaqMan probes. Gene expression levels were normalized to internal control *Hprt* and fold change increase in gene expression over naïve controls was calculated according to the ΔΔ cycle threshold (CT) method ^33^.

### Transcriptome analysis of isolated astrocytes

Astrocytes were isolated by magnetic microbeads from brain of Otud7b^fl/fl^ and GFAP-cre Otud7b^fl/fl^ mice, respectively, at day 7 p.i. Astrocytes isolated from naïve of Otud7b^fl/fl^ and GFAP-cre Otud7b^fl/fl^ mice were used as control. Total mRNA was isolated from purified astrocytes and RNA Library was created using a NEBNext® UltraTM II Directional RNA Library Prep Kit. The library was sequenced using a NovaSeq 6000 sequencer. The online platform DAVID Bioinformatics from the National Institutes of Health (NIH) was used for the KEGG pathway, molecular function, and biological process analyses.

### Pharmacological treatment

Pantethine was administered to mice by intraperitoneal injection to prevent microvesicle release during plasmodium infection. Pantethine was dissolved in sterile phosphate-buffered saline (PBS) and injected at the indicated dose of 15 mg per mouse starting day 2 post-infection (p.i.). Control animals received equivalent volumes of vehicle alone. Injections were performed under sterile conditions, and animals were monitored daily for signs of distress or adverse effects throughout the treatment period.

### Quantification and statistical analysis

Quantification of WB was performed using NIH ImageJ software. Statistical analysis and graphic design were performed using GraphPad Prism 10. The two-tailed Student’s t test was used to detect statistical differences in all experiments P values <0.05 (*) were considered statistically significant. All experiments were performed at least twice.

## Results

### OTUD7B in astrocytes exacerbates experimental cerebral malaria

To investigate the role of astrocytic OTUD7B during ECM, Otud7b^fl/fl^ control mice and GFAP-Cre Otud7b^fl/fl^ mice lacking OTUD7B in the astrocytes (Supplementary Fig 1A, B) were intraperitoneally injected with 1×10^6^ *Plasmodium berghei* ANKA (PbA) infected RBCs. The clinical disease progression was assessed using the Rapid Murine Coma and Behaviour Scale (RMCBS). Otud7b^fl/fl^ mice exhibited a significantly reduced RMCBS during disease progression compared with GFAP-Cre Otud7b^fl/fl^ mice, indicating that astrocytic OTUD7B exacerbates disease severity (Figure 1A). When OTUD7B-competent mice exhibited clinical signs of ECM (day 7 p.i.,), extravasation of Evans blue into the brain parenchyma was visible, illustrating disruption of BBB integrity. In contrast, the GFAP-Cre Otud7b^fl/fl^ mice showed no signs of disease and absence of Evans Blue extravasation in the brains of GFAP-Cre Otud7b^fl/fl^ mice demonstrated an intact BBB. (Figure 1B). Histological analysis at day 7 p.i. showed hemorrhagic lesions in the brains of Otud7b^fl/fl^ mice (Figure 1C). Immunohistochemical staining demonstrated increased activation of astrocytes and endothelial cells in Otud7b^fl/fl^ mice, compared to GFAP-Cre Otud7b^fl/fl^ mice as indicated by increased expression of GFAP, and CD31 (Figure 1D, E). Immunofluorescence analysis further revealed that astrocytes in Otud7b^fl/fl^ mice, identified by SOX2/9 and those located near blood vessels, exhibit a gliotic phenotype characterized by elevated GFAP and OTUD7B expression, along with a more gemistocytic morphology. In contrast, astrocytes in GFAP-Cre Otud7b^fl/fl^ mice, which lack OTUD7B, do not display gliosis and retain a normal appearance (Figure 1F). In addition, both immunohistochemistry (1G) and flow cytometry (Figure 1H, Supplementary Fig 1D) revealed a significant increase in CD8⁺ T cell infiltration in the brains of Otud7b^fl/fl^ mice compared to GFAP-Cre Otud7b^fl/fl^ mice at day 7 p.i. In contrast, Giemsa-stained thin blood films showed similar parasite loads in both Otud7b^fl/fl^ and GFAP-Cre Otud7b^fl/fl^ mice (Supplementary Fig 1C) indicating that astrocytic OTUD7B was not involved in pathogen control. Taken together these data indicate that OTUD7B in astrocytes promotes experimental cerebral malaria by disturbance of the neurovascular unit.

**Figure 1.**
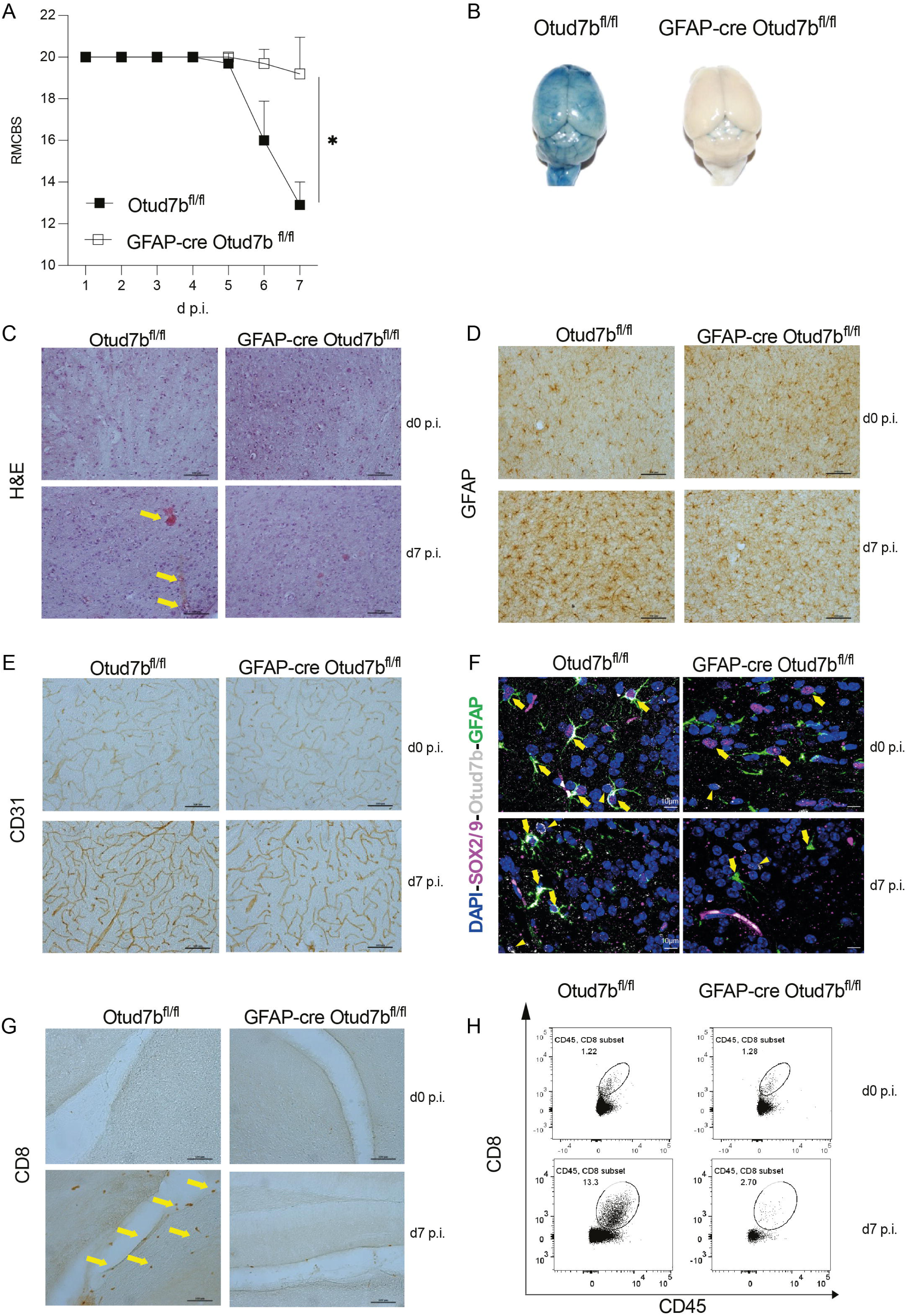
Astrocyte-specific deletion of OTUD7B attenuates experimental cerebral malaria. (A–I) Otud7b^fl/fl^ and GFAP-Cre Otud7b^fl/fl^ mice were injected intraperitoneally (i.p.) with 1 × 10^6^ *Plasmodium berghei* ANKA (PbA)-infected red blood cells (iRBCs). (A) Mice were monitored daily for neurological signs using the Rapid Murine Coma and Behavior Scale (RMCBS) until day 7 post-infection (p.i.). A score of 20 indicates a healthy mouse. Data represent one of two independent experiments (n = 10; Mann–Whitney U test, *p* < 0.05). (B) Representative Evans blue assay performed at day 7 p.i. Mice were injected intravenously (i.v.) with Evans blue dye and perfused with 0.9% saline. 1 hour later, brains were isolated and photographed. Two independent experiments were performed (n = 3 mice per group per experiment).(C) Hematoxylin and eosin staining showing hemorrhagic lesions throughout the brain at day 7 p.i. (D, E) Immunohistochemical staining showing activation of astrocytes (GFAP) and endothelial cells (CD31), respectively, at day 7 p.i. Scale bars, 100 μm. (F) Immunofluorescence staining demonstrating increased GFAP expression in brain sections of infected mice, with SOX2^+^/SOX9^+^/GFAP^+^ astrocytes surrounding hemorrhagic lesions. (G, H) Increased accumulation of CD8^+^ T cells in the brains of infected mice, assessed by immunohistochemistry (G) and flow cytometry (H) (Student’s *t*-test, *p* < 0.05).

### Astrocytic OTUD7B fosters proinflammatory gene expression in experimental cerebral malaria

To further characterize the regulation of neuroinflammation by astrocytic OTUD7B during ECM, brain tissue was harvested from infected Otud7b^fl/fl^ and GFAP-Cre Otud7b^fl/fl^ mice at day 7 p.i. and the expression of proinflammatory genes was analyzed by qRT-PCR. As shown in Figure 2A, the expression of the chemokines Cxcl9, Cxcl10, Cxcl11, the inflammatory cytokines IL-6 and TNF, and the cytotoxic effector molecules granzyme B (Gzmb) and perforin (Prf1) was higher in the brains of Otud7b^fl/fl^ compared with GFAP-Cre Otud7b^fl/fl^ mice.

**Figure 2.**
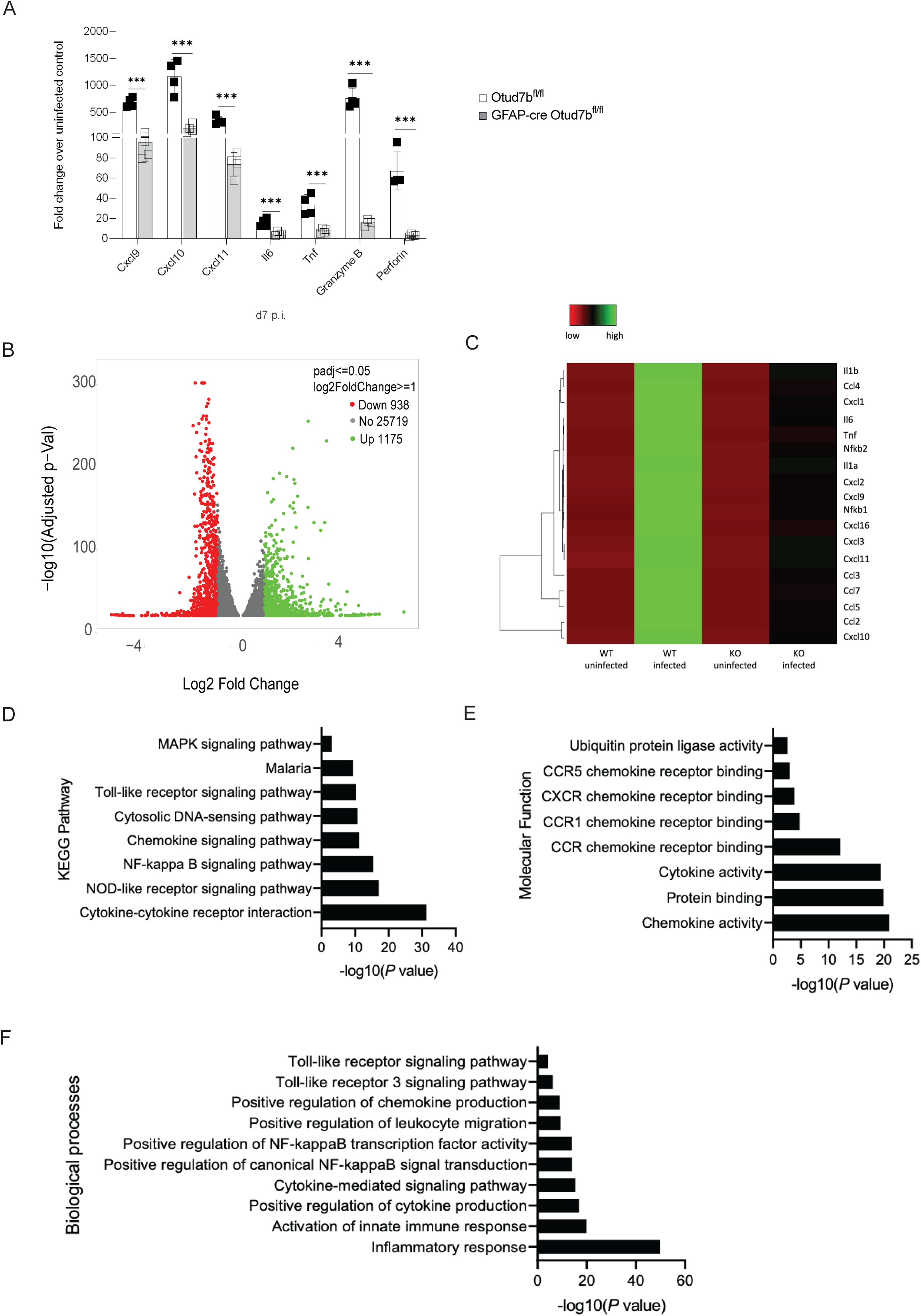
Impaired proinflammatory gene expression in astrocytes of GFAP-Cre Otud7b^fl/fl^ mice during experimental cerebral malaria. (A) Otud7b^fl/fl^ and GFAP-Cre Otud7b^fl/fl^ mice were injected i.p. with 1 × 10^6^ PbA-infected RBCs, and brains were harvested at day 7 p.i. Expression of chemokines (*Cxcl9*, *Cxcl10*, *Cxcl11*), cytokines (*Il6*, *Tnf*), and cytotoxic molecules (*Gzmb*, *Prf1*) was analyzed by quantitative RT-PCR and normalized to *Hprt*. Data are shown as fold change relative to uninfected controls (n = 4 per group). (B–F) Astrocytes were magnetically sorted from uninfected (day 0) and PbA-infected mice at day 7 p.i. and subjected to RNA sequencing. (B) Volcano plot showing differentially expressed genes between Otud7B sufficient and deficient astrocytes. (C) Heatmap of chemokine gene expression in astrocytes from uninfected and infected mice. (D–F) KEGG pathway (D), molecular function (E), and biological process (F) enrichment analyses were performed using DAVID bioinformatics tool (n = 3 per group).

To further study the OTUD7b-regulated astrocyte-intrinsic transcriptional changes, astrocytes were isolated from uninfected and infected Otud7b^fl/fl^ and GFAP-Cre Otud7b^fl/fl^ mice at day 7 p.i by magnetic sorting and RNA sequencing was performed. Differential gene expression analysis showed distinct transcriptional profiles between Otud7b^fl/fl^ and GFAP-Cre Otud7b^fl/fl^ mice with 1175 genes upregulated and 938 genes downregulated in OTUD7B-sufficient astrocytes compared to OTUD7B-deficient astrocytes as shown by the volcano plot (Figure 2B). Heatmap analysis showed enhanced induction of chemokine genes in astrocytes from infected Otud7b^fl/fl^ mice compared with astrocytes from GFAP-Cre Otud7b^fl/fl^ mice (Figure 2C). Functional enrichment analysis revealed increased activation of pathways related to immune signaling, chemokine activity, and inflammatory responses, in OTUD7B sufficient astrocytes as demonstrated by KEGG pathway, molecular function, and biological process analyses (Figure 2D–F). These data illustrate that OTUD7B promotes pro-inflammatory transcriptional activation of astrocytes during ECM.

CXCL10 is a biomarker for human cerebral malaria and is known to play an important role in the pathogenesis of ECM by recruiting pathogenic CD8 T cells to the brain ^34^. Our transcriptome data show increased production of CXCL10 by OTUD7b-competent astrocytes.

### Astrocytic OTUD7B enhances microvesicle-induced chemokine expression

Microvesicles derived from parasitized RBC play an important role in ECM pathogenesis ^21,22^. Since astrocytes do not come into direct contact with the infected RBCs, we hypothesized that uptake of microvesicles by astrocytes might activate pro-inflammatory signaling and that OTUD7B might control the magnitude of this signaling pathway. To prove this hypothesis, Otud7b^fl/fl^ and GFAP-Cre Otud7b^fl/fl^ mice were infected and the release of microvesicles was blocked using pantethine following a previously published study ^35^. To monitor the efficiency of microvesicle inhibition, the microvesicles, characterized as Annexin V⁺ and based on size and side scatter characteristics, were quantified in serum of infected mice by flow cytometry (Figure 3A). PbA infection resulted in a similar increase in microvesicle numbers between Otud7b^fl/fl^ and GFAP-Cre Otud7b^fl/fl^ mice and pantethine treatment equally reduced the microvesicle numbers in both genotypes (Figure 3A). Inhibition of microvesicles significantly (p > 0.001) improved the disease outcome of Otud7b^fl/fl^ and increased their RMCBS as compared PBS-treated Otud7b^fl/fl^ mice (Fig. 3B). In addition, pantethine treatment prevented Evans Blue extravasation in infected Otud7b^fl/fl^ mice indicating prevention of BBB disruption in these animals (Fig. 3C) and also significantly reduced cerebral expression of the chemokines Cxcl9, Cxcl10, and Cxcl11 (Figure 3D). In contrast, pantethine did not impact on disease activity, BBB integrity and chemokine production in GFAP-Cre Otud7b^fl/fl^ mice (Figure 3B, C, D) indicating that astrocytic OTUD7B was required to induce microvesicle induced proinflammatory signaling and disease in ECM.

**Figure 3.**
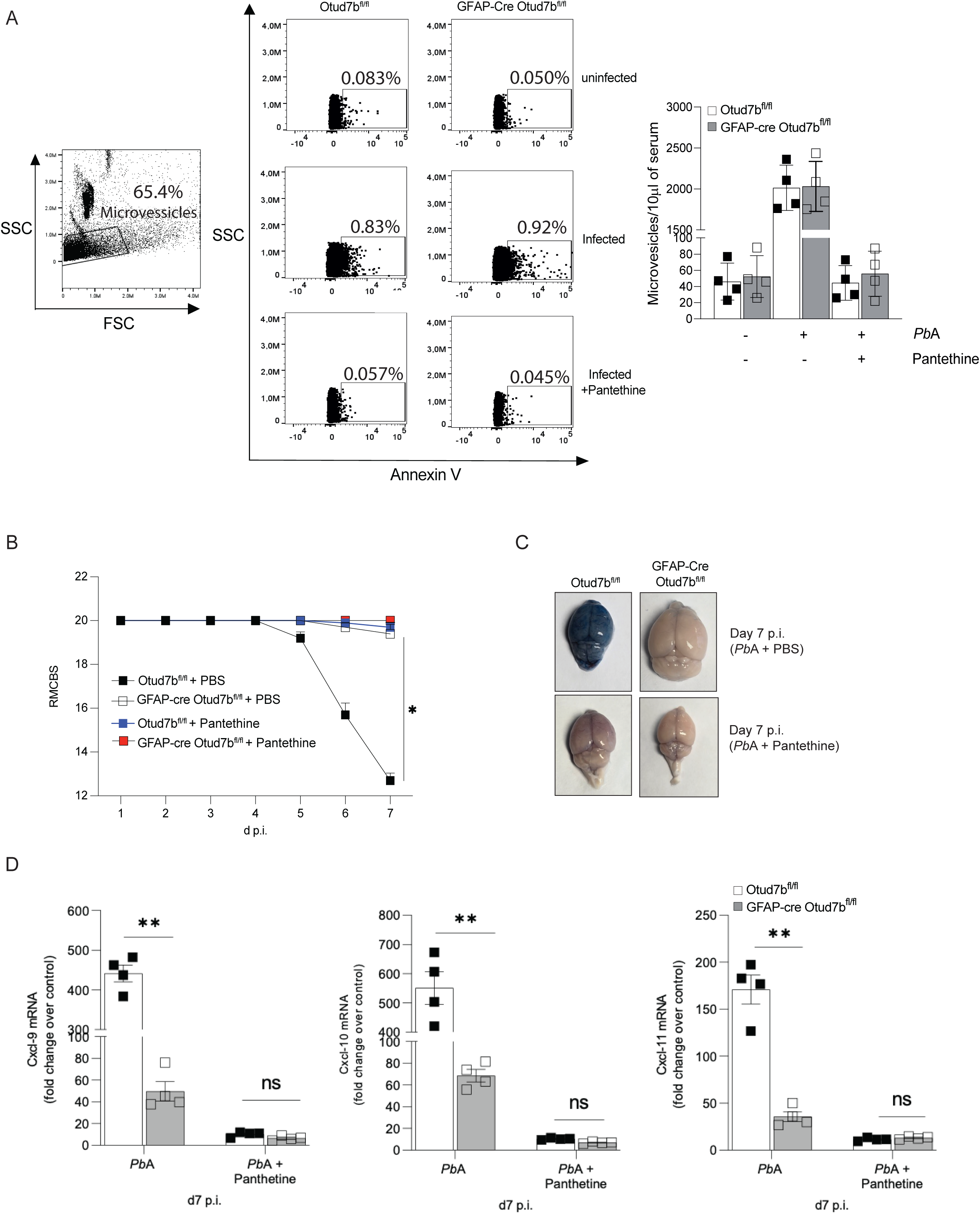
Pantethine treatment reduces microvesicle shedding and protects mice from cerebral malaria. (A–E) Otud7b^fl/fl^ and GFAP-Cre Otud7b^fl/fl^ mice were injected i.p. with 1 × 10^6^ PbA-infected RBCs. Mice received intraperitoneal injections of pantethine (15 mg in saline) everyday starting day 2 p.i. (A) Representative flow cytometry dot plot showing Annexin V^+^ microvesicles identified based on size (side scatter) relative to calibration microbeads and the absolute number of Annexin V^+^ microvesicles in 10 μl serum from untreated and pantethine-treated mice. (B) RMCBS neurological scores monitored daily until day 7 p.i. (C) Representative Evans blue assay at day 7 p.i. (D) Brain expression levels of *Cxcl9*, *Cxcl10*, and *Cxcl11* were measured by qRT-PCR and normalized to *Hprt*.

### OTUD7B promotes TRAF3- and TRAF6-dependent NF-κB and MAP kinase signaling in astrocytes

To delineate the signal transduction pathways regulated by OTUD7B in astrocytes, primary astrocytes isolated from Otud7b^fl/fl^ and GFAP-Cre Otud7b^fl/fl^ mice were stimulated with serum-derived microvesicles. Infection induced microvesicles were isolated from the serum of mice by standard protocols ^36^ and are characterized as Annexin V^+^. Representative picture showing ACSA-2 positive astrocytes stimulated with Annexin V^+^ microvesicles (Supplementary Fig. 1E). At 6 and 24 h post stimulation with microvesicles, OTUD7B-sufficient astrocytes displayed significantly enhanced induction of proinflammatory genes, including Cxcl9, Cxcl10, Cxcl11, Ccl2, Il6, and Tnf, compared with OTUD7B-deficient astrocytes (Figure 4A). Western blot analysis showed that the protein levels of MYD88, IRAK1/4 and TAK1 did not differ between the groups (Figure 4B). However, the levels of the adaptor molecules TRAF3 and TRAF6 remained stable over time in OTUD7B-sufficient astrocytes after microvesicle stimulation but declined in OTUD7B-deficient astrocytes (Figure 4B). This is associated with enhanced downstream activation of NF-κB (p-p65) and MAP kinase (p-p38) pathways in OTUD7B-sufficient as compared to OTUD7B-deficient astrocytes (Figure 4B). These data suggest that OTUD7B determined the magnitude of NF-κB and MAP kinase signaling by regulating TRAF3/TRF6 stability.

**Figure 4.**
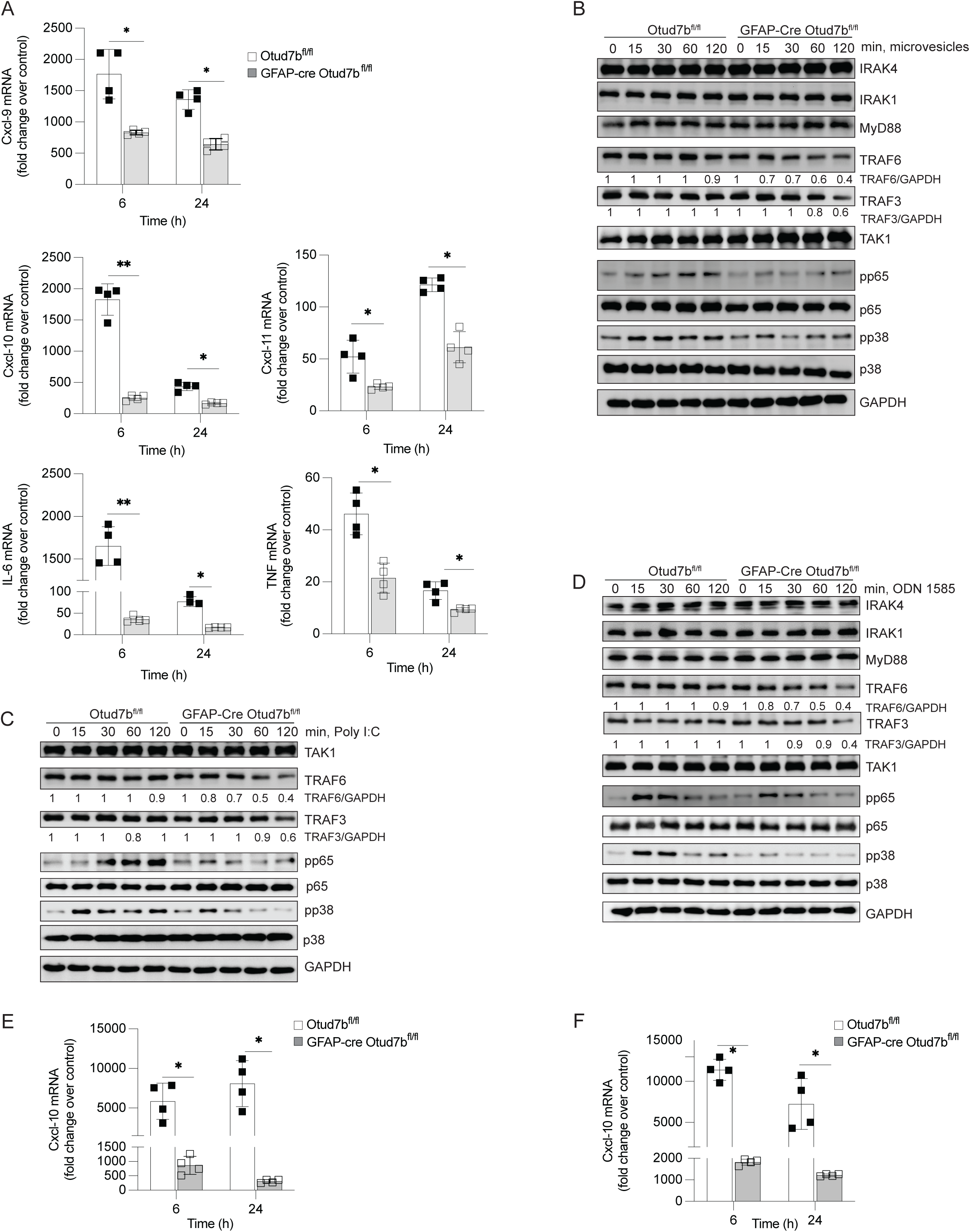
OTUD7B promotes proinflammatory responses in astrocytes upon microvesicle stimulation. (A,B) Primary astrocytes were isolated from P0–P1 pups of Otud7b^fl/fl^ and GFAP-Cre Otud7b^fl/fl^ mice and stimulated with serum-derived microvesicles. (A) Relative mRNA expression of *Cxcl1*, *Cxcl10*, *Cxcl11*, *Ccl2*, *Il6*, and *Tnf* was measured by qRT-PCR (n = 4). Data represent fold change relative to unstimulated controls (mean ± SEM; Student’s *t*-test, *p* < 0.05, \**p* < 0.01, \*\**p* < 0.001). (C–F) Protein lysates were analyzed by immunoblotting following stimulation with microvesicles (C), Poly I:C (10 µg/ml) (D), or ODN1585 (10 µg/ml) (E). Representative blots from three independent experiments are shown. (E,F) Relative *Cxcl10* mRNA expression following Poly I:C (E) or ODN1585 (F) stimulation was quantified by qRT-PCR (n = 4).

In agreement, following stimulation with the TLR3 ligand Poly I:C (Figure 4C, E) and the TLR9 ligand ODN1585 (Figure 4D, F), respectively, a similar reduction in levels of TRAF3/TRAF6 (Figure 4C, D), diminished activation of NF-κB and MAP kinase (Figure 4C, D) and production of Cxcl10 (Figure 4E, F) was observed in OTUD7B-deficient astrocyte.

Next, we validated the hypothesis that microvesicles of infected mice induce and OTUD7B regulates the TRAF3/TRAF6-dependent activation of NF-κB and MAP kinase and that NF-κB and MAP kinase mediate chemokine production in astrocytes. First, we silenced TRAF3 and TRAF6, respectively, using siRNAs in OTUD7B-competent and -deficient astrocytes, scrambled siRNA was used as a control (Figure 5A, B). TRAF6 siRNA but not TRAF3 siRNA treatment prevented p65 and p38 phosphorylation in both genotypes (Figure 5A). In addition, TRAF6 but not TRAF3 siRNA inhibited Cxcl10 mRNA expression (Figure 5B) indicating that TRAF6 but not TRAF3 is critical for microvesicle-induced NF-κB and MAP kinase activation and Cxcl10 production. To further delineate that microvesicle-induced NF-κB and/or MAP kinase activation drives OTUD7B-regulated Cxcl10 mRNA production, we treated microvesicle activated OTUD7B-competent and -deficient astrocytes with an IKK/NF-κB (p65) and p38 MAP kinase inhibitor, which both prevented activation of the respective signaling molecules (Figure 5C). Inhibition of either IKK or p38 reduced expression of Cxcl10 in OTUD7B-sufficient astrocytes but still Cxcl10 production was lower in OTUD7B-deficient astrocytes (Figure 5D). However, combined treatment with both inhibitors diminished Cxcll0 mRNA to equally low levels in the two genotypes. Collectively, these data show that (i) microvesicles derived from infected mice induce TRAF6-dependent activation of NF-κB (p65) and p38 MAP kinase, which both induce Cxcl10 production in astrocytes and (ii). OTUD7b augments this microvesicle-induced pro-inflammatory astrocyte activity.

**Figure 5.**
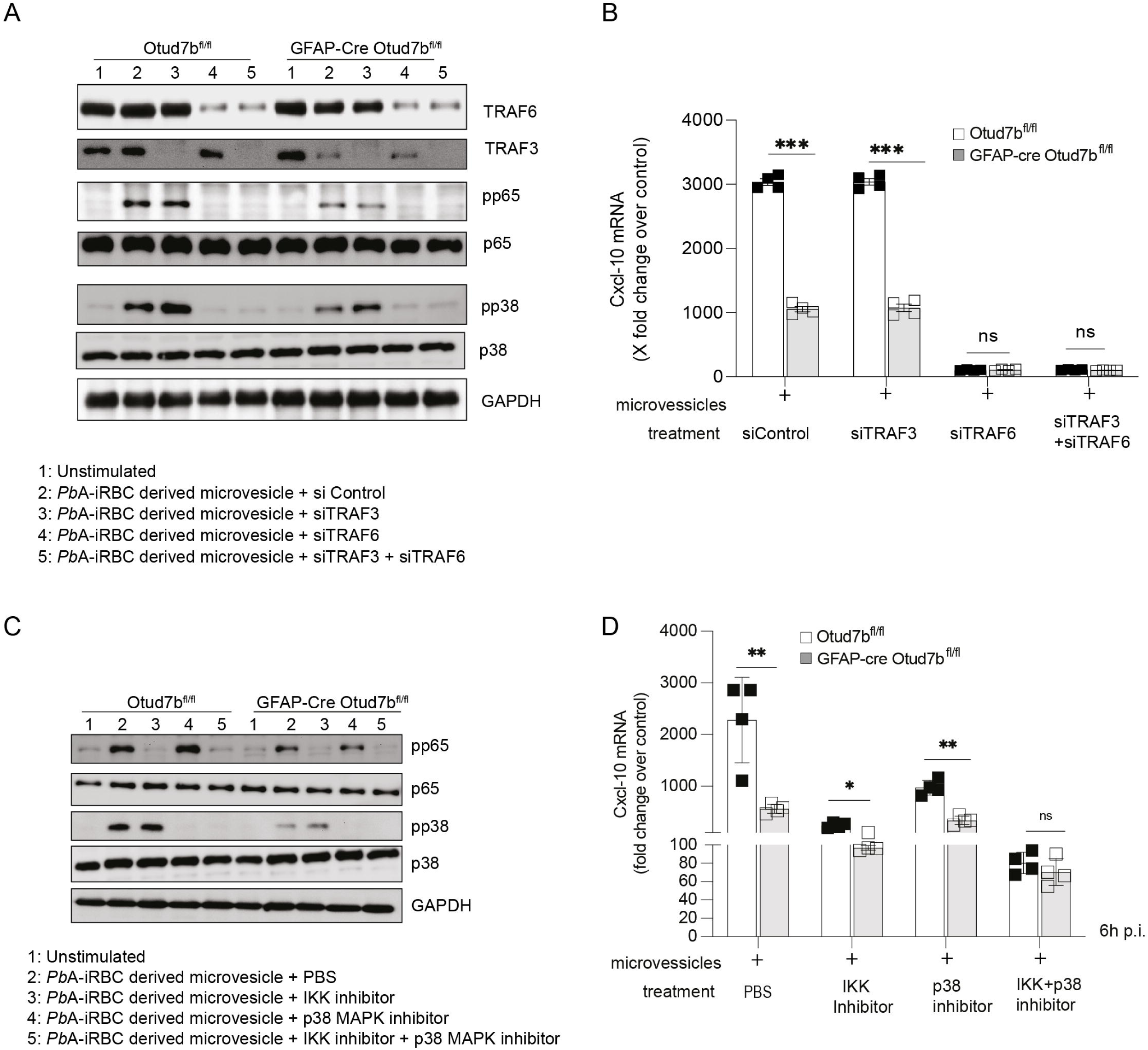
OTUD7B promotes TRAF3- and TRAF6-mediated NF-κB and MAP kinase–dependent proinflammatory responses in astrocytes. (A, B) TRAF3 and TRAF6 were silenced individually or in combination using siRNA in OTUD7B-sufficient and OTUD7B-deficient astrocytes. Cells were stimulated with microvesicles for 6 hours. (A) Immunoblot analysis of TRAF3, TRAF6, phosphorylated and total p65, and phosphorylated and total p38. GAPDH served as loading control. (B) *Cxcl10* mRNA levels were measured by qRT-PCR following stimulation with PbA-derived microvesicles. (C, D) Primary astrocytes from P0–P1 Otud7b^fl/fl^ and GFAP-Cre Otud7b^fl/fl^ mice were pretreated for 2 hours with either the NF-κB inhibitor IKK Inhibitor VII or the p38 MAP kinase inhibitor SB203580, followed by stimulation with serum-derived microvesicles. (C) Protein lysates were analyzed by immunoblotting using the indicated antibodies. (D) Relative *Cxcl10* mRNA expression was quantified by qRT-PCR (n = 4). Data are presented as fold change relative to unstimulated controls (mean ± SEM).

### OTUD7B removes K48-linked ubiquitin chains from TRAF3 and TRAF6

To determine how OTUD7B regulates TRAF6 and TRAF3 function, we first studied whether OTUD7B interacts with both molecules. Primary astrocytes isolated from Otud7b^fl/fl^ and GFAP-Cre Otud7b^fl/fl^ mice were stimulated either with microvesicles (Figure 6A, D, G), Poly I:C (Figure 6B, E, H), or ODN1585 (Figure 6C, F, I) respectively. Co-immunoprecipitation experiments showed an interaction of OTUD7B with both TRAF3 and TRAF6 (Figure 6A–C). Furthermore, immunoprecipitation of TRAF3 (Figure 6D–F) and TRAF6 (Figure 6G–I) showed increased K48-linked polyubiquitin chains in OTUD7b-deficient astrocytes, indicating that OTUD7B stabilizes TRAF3 and TRAF6 by cleaving K48-linked ubiquitin chains.

**Figure 6.**
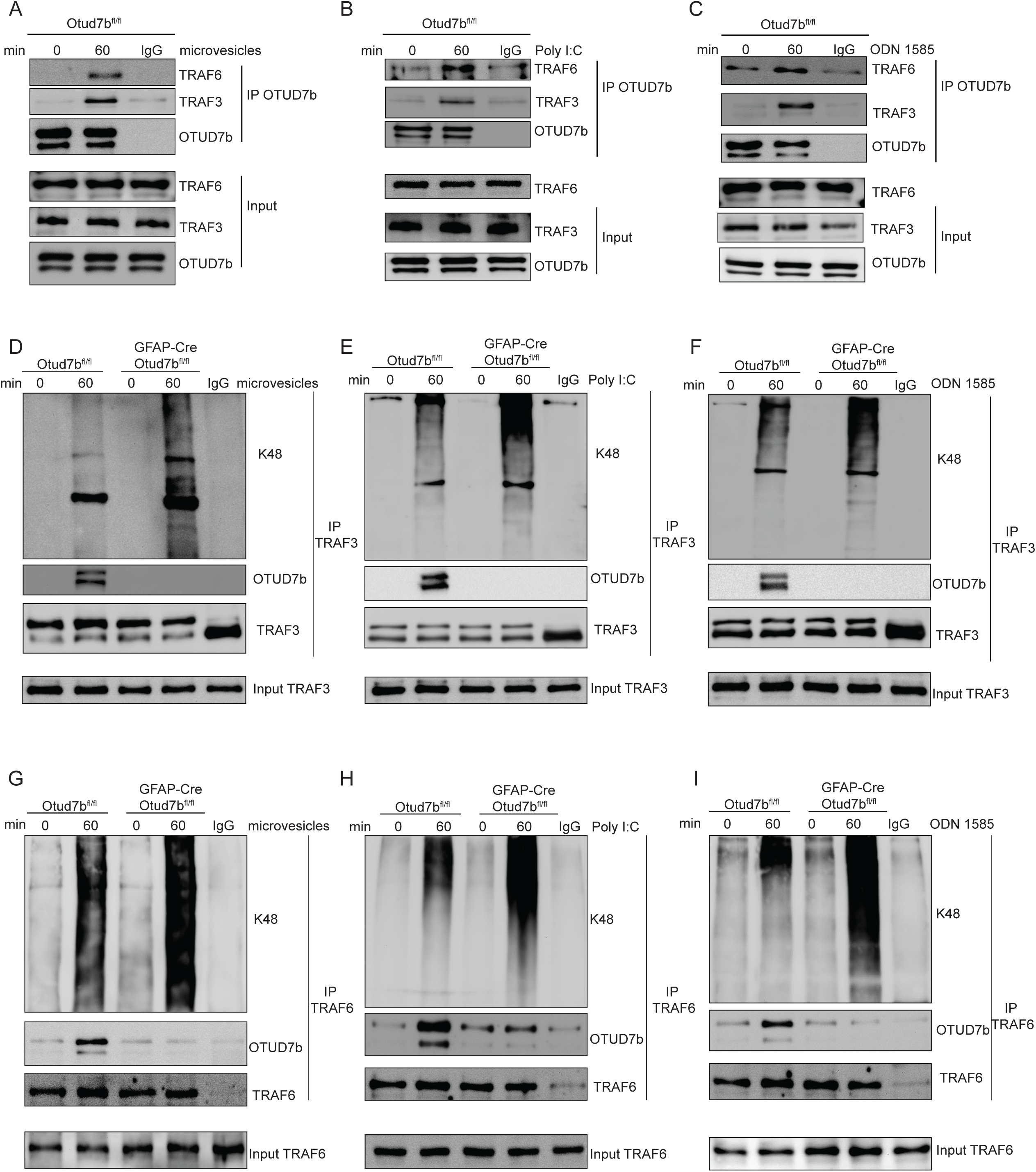
OTUD7B removes K48-linked ubiquitin chains from TRAF3 and TRAF6. (A–I) OTUD7B-sufficient and OTUD7B-deficient astrocytes were stimulated with microvesicles (A, D, G), Poly I:C (B, E, H), or ODN1585 (C, F, I). Cells were harvested at 0 and 60 minutes post-stimulation. (A–C) OTUD7b was immunoprecipitated, and its interaction with TRAF3 and TRAF6 was analyzed by immunoblotting. (D–F) TRAF3 was immunoprecipitated, and association with OTUD7B and K48-linked ubiquitination levels were assessed. (G–I) TRAF6 was immunoprecipitated, and interaction with OTUD7B and K48-linked ubiquitination status were determined.

### Therapeutic inhibition of cerebral Otud7b or CXCL10 protects mice from cerebral malaria

To evaluate whether OTUD7B could serve as a therapeutic target to treat ECM, Otud7b^fl/fl^ mice were treated intranasally with *Otud7b* siRNA at day 5 p.i. infection. At this stage of infection, Otud7b^fl/fl^ mice exhibit first neurological symptoms, whereas GFAP-Cre Otud7b^fl/fl^ mice do not develop clinical neurological symptoms. The efficiency of the knockdown was validated by WB (Figure 7A). Data show high OTUD7B protein expression in olfactory bulb, cortex and brain stem of Otud7b^fl/fl^ mice (Figure 7A). GFAP-Cre Otud7b^fl/fl^ mice had lower expression of OTUD7B in these brain regions, presumably due to absence of OTUD7B protein in astrocytes. Treatment with *Otud7b* siRNA completely abolished OTUD7B protein expression at day 7 p.i. (Figure 7A) Inhibition of cerebral OTUD7B significantly improved, i.e. increased, the neurological RMCBS scores of *Otud7b* siRNA treated mice compared with control-treated Otud7b^fl/fl^ mice (Figure 7B). Evans blue assays showed reduced vascular leakage in siRNA-treated mice Otud7b^fl/fl^ mice, which was identical to GFAP-Cre Otud7b^fl/fl^ mice (Figure 7C). Furthermore, cerebral expression of Cxcl10 mRNA was also significantly reduced following *Otud7b* knockdown in Otud7b^fl/fl^ mice (Figure 7D). To validate the importance of cerebral CXCL10 in ECM, Otud7b^fl/fl^ mice were treated intranasally with *cxcl10* siRNA at day 5 p.i. infection. The reduction in *cxcl10* mRNA was validated by qRT-PCR (Figure 7E). Inhibition of *cxcl10* improved the clinical scores of siRNA-treated Otud7b^fl/fl^ mice (Figure 7F) and prevented the BBB disruption (Figure 7G) which was comparable to GFAP-Cre Otud7b^fl/fl^ mice. Collectively Astrocytic OTUD7B fosters microvesicle-driven TRAF3/TRAF6 activation, sustaining NF-κB and MAPK signaling leading to neuroinflammation and enhanced CXCL10 production. This recruits pathogenic CD8⁺ T cells to the brain, which contribute to BBB disruption and severe neuroinflammation in cerebral malaria (Figure 8A, B).

**Figure 7.**
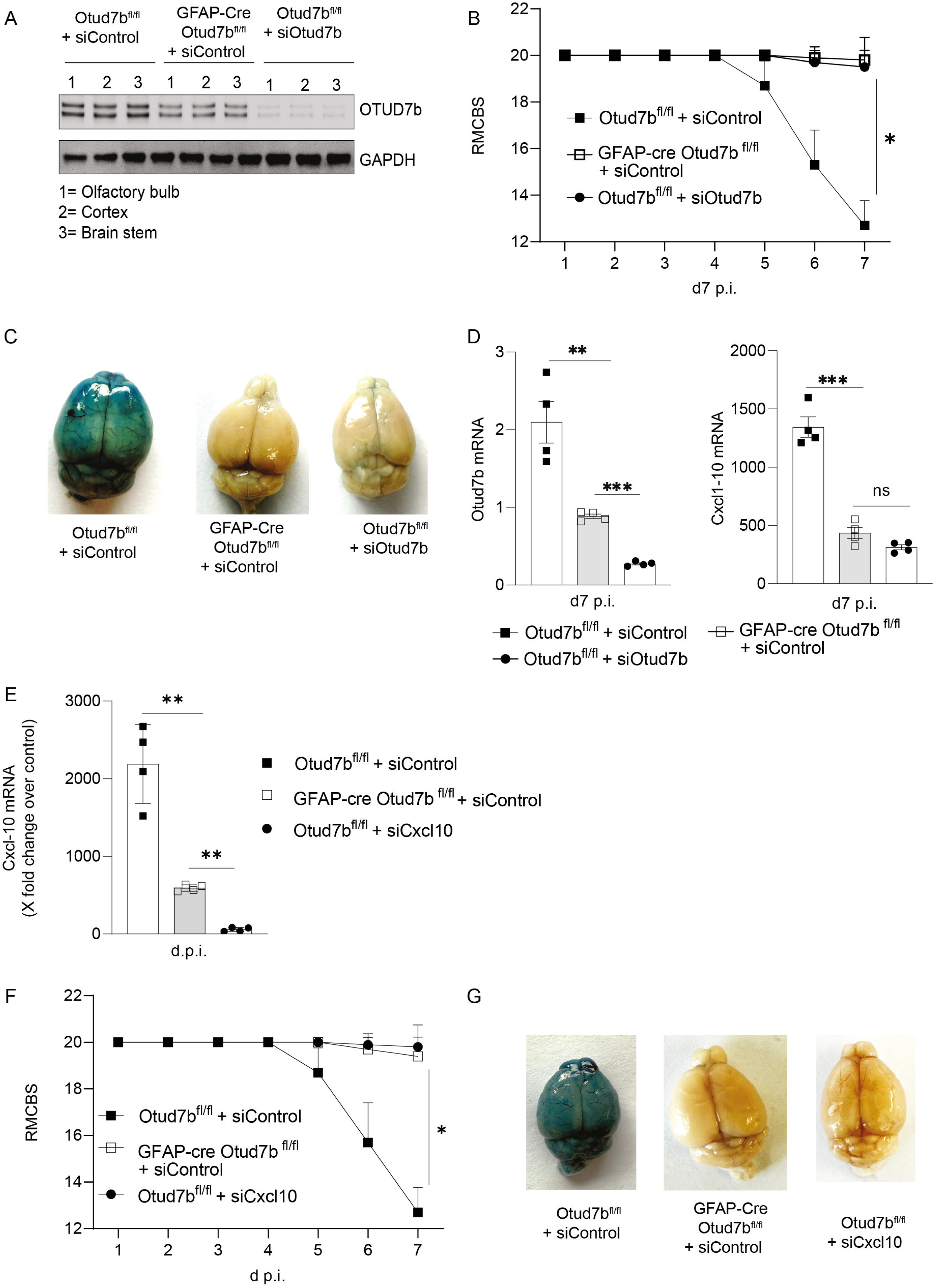
Therapeutic inhibition of cerebral *Cxcl10* or *Otud7b* protects mice from cerebral malaria. (A-G) Otud7b^fl/fl^ and GFAP-Cre Otud7b^fl/fl^ mice were injected i.p. with 1 × 10^6^ PbA-infected RBCs. (A-D) Mice received intranasal administration of *Otud7b* siRNA or scrambled control siRNA on day 5 p.i. (A) Representative western blot showing knockdown of OTUD7B in different brain regions after siRNA treatment (B) Neurological scores (RMCBS) were monitored daily until day 7 p.i. (C) Representative Evans blue assay at day 7 p.i. (D) Brain *Otud7b* and *Cxcl10* expression was analyzed by qRT-PCR and normalized to *Hprt*. Mice either received intranasal Cxcl10 siRNA or scrambled control on day 5 p.i. (E) The knock down of cerebral *Cxcl10* mRNA expression was monitored by qRT-PCR. (F) Neurological scores (RMCBS) were monitored daily until day 7 p.i. (G) Representative Evans blue assay. (H)

**Figure 8.**
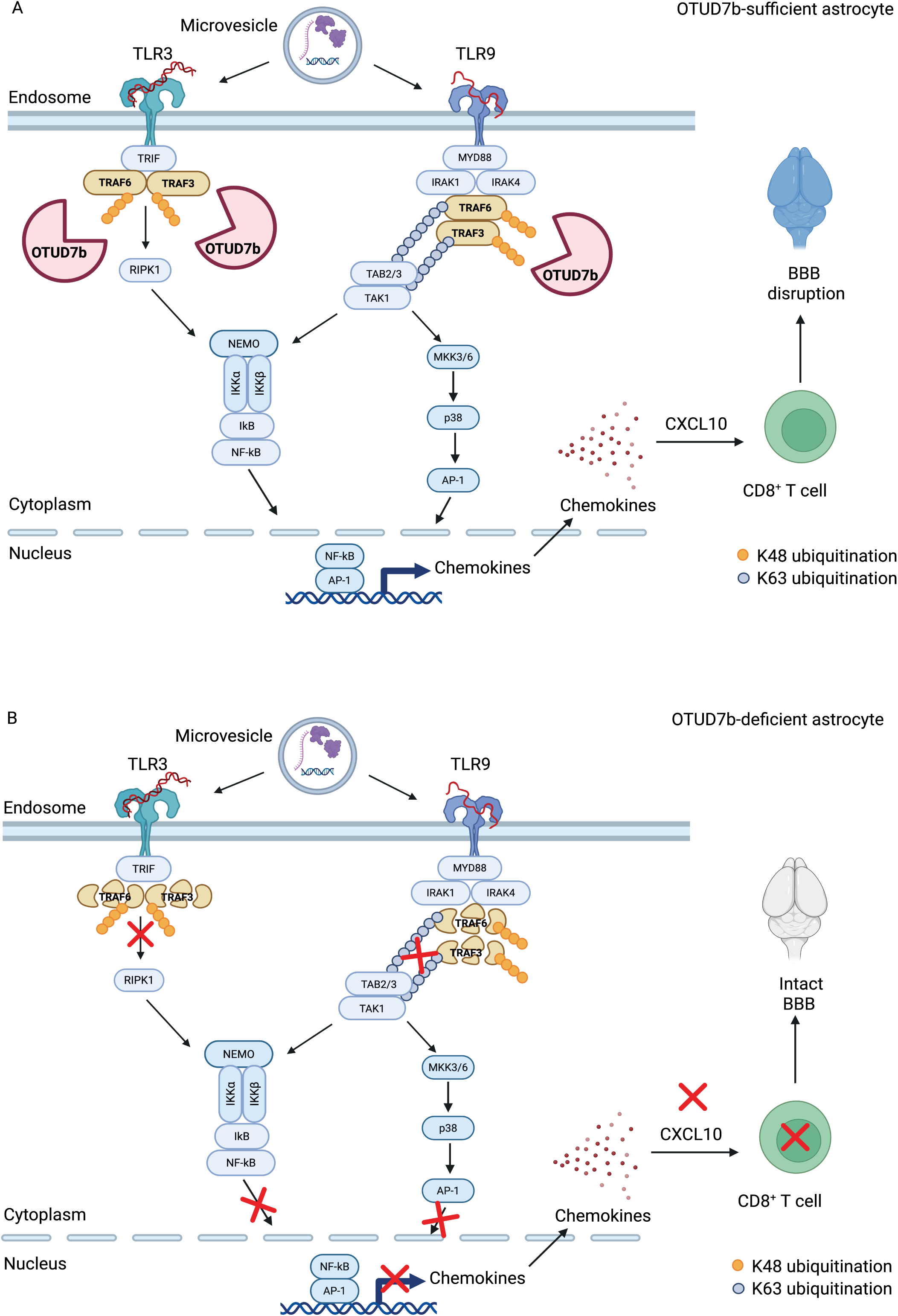
Astrocytic OTUD7B fosters microvesicle induced cerebral malaria. A) During cerebral malaria, OTUD7B fosters microvesicle-induced activation of NF-κB and MAPK signaling by cleaving K48-linked ubiquitin chains from TRAF3/TRAF6 and stabilizing these molecules. This results in elevated production of the chemokine CXCL10, which leads to increased recruitment of pathogenic CD8⁺ T cells and subsequent disruption of BBB and severe neuroinflammation. B) Deletion of OTUD7B in astrocytes results in degradation of TRAF3/TRAF6, which limits the activation of the NF-κB and MAPK pathways and controls the production of CXCL10. This restricts the recruitment of CD8⁺ T cells to the brain, preserves BBB integrity, and protects against cerebral malaria. The figure was created using BioRender (Gopala Nishanth, 2025; https://biorender.com/rin8c1c)

## Discussion

This study identifies astrocytes as key players determining the outcome of ECM by modulation of the proinflammatory responses and neurovascular pathology. This largely extents the concept that intracerebral sequestration of parasitized erythrocytes, intracerebral accumulation of cytotoxic CD8^+^ T cells, interaction of T cells with endothelial interaction and endothelial dysfunction are the main drivers of ECM ^11,37,38^. Although astrocytes based on their anatomic location in the BBB do not come into direct contact with infected erythrocytes, the identification that astrocytes are directly activated by microvesicles carrying parasitic cargo provides a mechanistic explanation how their OTUD7B-dependent chemokine production can be induced and participate in the intricate interplay of brain resident cells, parasitized erythrocytes and immune cells in ECM.

Our transcriptome analysis of ex vivo–isolated astrocytes showed that OTUD7B strongly augments the production of pro-inflammatory chemokines and cytokines during ECM. These cytokines and chemokines play an important role in the pathogenesis of ECM ^39^. Cytokines IFN- γ, IL-1β, and IL-6 induce the expression of cell adhesion molecules (ICAM-1, VCAM-1) on endothelial cells, which contributes to the sequestration of parasitized erythrocytes in cerebral blood vessels. In addition, the CXCR3 ligands CXCL9 and CXCL10 mediate the recruitment of pathogenic CD8⁺ T cells to the brain, stabilize the interaction of these CD8^+^ T cells with endothelial cells expressing cell adhesion molecules and, thereby, determine the severity of ECM ^34,40,41^. Our study provides evidence that the OTUD7B-dependent production of CXCL10 by astrocytes is a critical step in the intracerebral accumulation of CD8^+^ T cells, breakdown of the BBB and development of ECM.

The functional relevance of OTUD7B-dependent production of CXCL10 by astrocytes is illustrated by the prevention of severe ECM by *Otud7b* siRNA treatment, which significantly reduced CXCL10 production, and by *Cxcl10* siRNA treatment, which also prevented development of severe ECM in GFAP-Cre Otud7b^fl/fl^ mice. Importantly, intranasal treatment with both *Otud7b* and *Cxl10* siRNA, respectively, at the onset of neurological signs of ECM (day 5 p.i.) was sufficient to prevent progressions of lethal ECM. Thus, therapeutic inhibition of inflammatory checkpoints in the brain can ameliorate ECM trajectory even after the onset of the disease. These data along with our in vivo observations of diminished cerebral inflammation in the absence of astrocytic OTUD7B provide a proof-of-principle that targeting astrocytes, which are the major inducers of cerebral inflammation could be an attractive adjunct therapeutic strategy to reduce neuropathology in CM.

Mechanistically, we uncovered that stimulation of astrocytes with microvesicles isolated from Plasmodium-infected mice induced OTUD7B-dependent activation of canonical NF-κB and p38 MAP kinases. Microvesicles are released from infected erythrocytes but also endothelial cells, platelets and leukocytes during malaria and contain parasitic proteins, RNA, DNA, lipids, and also TLR3 and TLR9 activating ligands ^42–44^. They have been proposed to promote disease severity ^21,22^. Our study shows that microvesicles can trigger inflammatory signaling in astrocytes and inhibition of microvesicle release using pantethine confers protection from ECM. This indicates that the systemic infection contributes to CNS pathology through microvesicle-mediated signaling, which in turn triggers production of chemokines and cerebral recruitment of leukocytes by astrocytes. Upon stimulation of astrocytes with microvesicles, or with TLR3 and TLR9 agonists, OTUD7B interacts with both TRAF3 and TRAF6, specifically cleave K48-linked polyubiquitin chains from these signaling adaptors. The interaction between OTUD7B and TRAF3 aligns with findings from Hu et al. (2013), who reported a similar interaction in B cells upon LTβR stimulation ^27^. The authors show that OTUD7B cleaves K48-linked polyubiquitin chains from TRAF3, thereby suppressing aberrant activation of the non-canonical NF-κB pathway ^27^. However, our study reveals a different function of this interaction in astrocytes. Upon microvesicle-induced stimulation, OTUD7B-mediated deubiquitination of TRAF3 sustains canonical NF-κB activation and p38 MAPK signaling, ultimately driving the production of the chemokine CXCL10. Regarding TRAF6, our findings extend previous work by Luong et al. (2013), who showed that OTUD7B cleaves K63-linked polyubiquitin chains from TRAF6 in HUVECs under hypoxia and thereby inhibits NF-κB mediated inflammatory responses ^45^. In contrast, in astrocytes, the removal of K48 linked polyubiquitin chains from TRAF6 stabilized TRAF6 and lead to the sustained activation of canonical NF-κB and p38 MAPK mediated inflammation. This suggests that regulation of TRAF3 and TRAF6 by OTUD7B critically depends upon the ubiquitin linkage, the cell type and the underlying stimulus.

Inhibition experiments in microvesicle-activated OTUD7B-expressing astrocytes revealed that TRAF6 as compared to TRAF3 had a superior role in the activation of canonical NF-κB and p38 MAP kinase and CXCL10 production, and that both NF-κB and p38 MAP kinase synergistically induced CXCL10 production. Although these data imply that the in vivo effects of OTUD7B silencing are caused by the inhibition in astrocytes, further studies specifically targeting OTUD7B in astrocytes are needed.

Notably our previous study showed that astrocytic OTUD7B restricts rather than augments neuroinflammation in a murine model of CNS autoimmunity, i.e., experimental autoimmune encephalomyelitis (EAE)^10^. In this model OTUD7B impaired TNF-induced neuroinflammation by limiting RIPK1 activation and preventing the degradation of the cytoskeletal protein GFAP ^32^. Since TNF neutralization did not improve the course of CM ^7^, the different inflammatory milieu and astrocyte-activating stimuli, i.e TNF in EAE and microvesicles with parasitic cargo in ECM, may account for the different function of astrocytic OTUD7B in these two models of neuroinflammation. This assumption is supported by studies which showed that OTUD7B dampens TNF- and non-canonical NF-κB driven inflammation ^27,29^. Moreover, our study on OTUD7B in dendritic cells indicates that in addition to the inflammatory milieu, the underlying cell type plays an important role determining the function of OTUD7B. In early stages of ECM, OTUD7B prevents TNF-induced apoptosis of dendritic cells by deubiquitination of the E3 ligase TRAF2 ^31^. Collectively these findings indicate that the functional role of OTUD7B depends both on the respective cell type and the inflammatory milieu.

In summary, our study demonstrates for the first time that astrocytes directly regulate the outcome of experimental cerebral malaria and identifies OTUD7B as a disease and cell type dependent driver of astrocyte inflammation.

## Acknowledgments

The authors thank Rituparna Bhattacharjee, Birgit Brennecke, Kerstin Ellrott, and Nadja Schlüter for expert technical assistance. This work was supported by a grant from the DFG CRC 854, project A30N to GN and by DFG EXC 2155 “RESIST” (Project ID39087428 to DS).

## Author contributions

GN conceptualized and supervised the experiments. KH, JS, and GN performed experiments and analyzed data. KH, RB, DS and GN interpreted the data. KH, DS, and GN wrote the manuscript.

## Supplementary Figure

**Supplementary Figure 1. Astrocytic OTUD7B contributes to cerebral malaria pathogenesis**

(A) Immunofluorescence staining of brain sections, arrows show Sox2/9 and GFAP positive astrocytes express high levels of OTUD7b in Otud7b^fl/fl^ mice while the astrocytes of GFAP-Cre Otud7b^fl/fl^ mice showed no expression of OTUD7B. DAPI was used as nuclear counterstain. (B) Immunoblot analysis of ex vivo–isolated astrocytes confirming OTUD7B deletion in GFAP-Cre Otud7b^fl/fl^ mice, (C) Peripheral parasitemia was monitored daily by microscopy of Giemsa-stained thin blood smears (n = 10 per group; two independent experiments). (D) Absolute numbers of CD45^+^CD8^+^ T cells in the brains of uninfected and PbA-infected mice at day 7 p.i. (E) Representative picture showing uptake of Annexin V^+^ microvesicle, stained green by ACSA-2^+^ astrocytes, stained in red. The nucleus was stained using DAPI

