## Supplementary figures and images for "Deletion of OTUD7B in astrocytes protects against cerebral malaria by inhibiting microvesicle-induced TRAF3/TRAF6-mediated neuroinflammation"

### Supplementary Figure 1

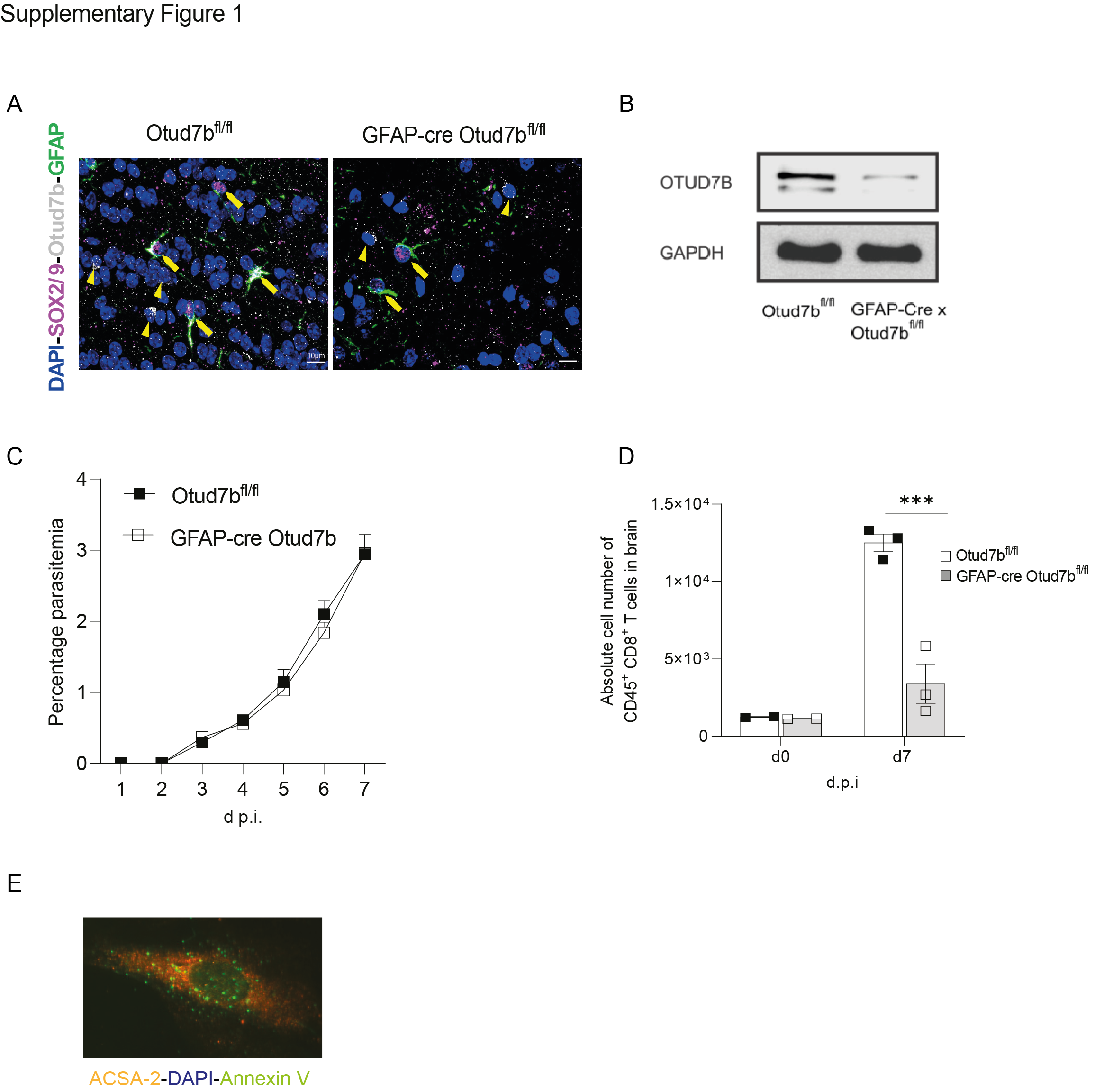
